# Systematic identification of factors bound to isolated metaphase ESC chromosomes reveals a role for chromatin repressors in compaction

**DOI:** 10.1101/750067

**Authors:** Dounia Djeghloul, Anne-Céline Kohler, Bhavik Patel, Holger Kramer, Nicolas Veland, Chad Whilding, Andrew Dimond, James Elliott, Amelie Feytout, Tanmay A.M. Bharat, Abul K. Tarafder, Jan Löwe, Bee L. Ng, Ya Guo, Karen Brown, Jacky Guy, Matthias Merkenschlager, Amanda G. Fisher

## Abstract

Epigenetic information is transmitted from mother to daughter cells through mitosis. To identify trans-acting factors and cis-acting elements that might be important for conveying epigenetic memory through cell division, we isolated native (unfixed) chromosomes from metaphase-arrested cells using flow cytometry and performed LC-MS/MS to determine the repertoire of chromosome-bound proteins. Quantitative proteomic comparisons between metaphase-arrested cell lysates and chromosome-sorted samples revealed a cohort of proteins that were significantly enriched on mitotic ESC chromosomes. These include pluripotency-associated transcription factors, repressive chromatin-modifiers (such as PRC2 and DNA methyl-transferases) and proteins governing chromosome architecture. We showed that deletion of PRC2, DNMT1/3a/3b or Mecp2 provoked an increase in the size of individual mitotic chromosomes consistent with de-condensation, as did experimental cleavage of cohesin complexes. These data provide a comprehensive inventory of chromosome-bound factors in pluripotent stem cells at mitosis and reveal an unexpected role for chromatin repressor complexes in preserving mitotic chromosome compaction.

## Introduction

Cell division requires genetic and epigenetic information to be accurately duplicated and conveyed to daughter cells. This process relies upon DNA synthesis at S-phase and the subsequent segregation of copies during mitosis (M) phase. In recent years, huge progress has been made in understanding not only how chromosomal DNA is copied and segregated, but also how epigenetic information is transmitted through the cell cycle. For example, we know that DNA methylation is reinstated during S-phase through the activity of DNMT1, an enzyme that re-establishes methylation at hemi-methylated CpG sites^1, 2^. It has also been proposed that during S-phase the epigenetic modifier Polycomb Repressor Complex 2 (PRC2) can both methylate histone H3 at lysine 27 (through Ezh2) and recognise this mark (through Eed)^3^, ensuring that histone H3K27 methylation is copied at newly synthesised DNA strands^4^. Understanding how epigenetic information is then transmitted through mitosis, as newly replicated genomes condense and segregate, is the subject of intense investigation but remains only partially understood. Progressive activation of CyclinB1-Cdk1 promotes chromosome condensation^5^ so that visibly discrete individual mitotic chromosomes appear at the mitotic spindle ahead of breakdown of the nuclear envelope. The physical basis of this chromosome compaction, as cells move from interphase into metaphase, remains somewhat controversial with evidence supporting at least two different models. The hierarchical model describes an ordered folding of DNA into consecutive higher-order structures, beginning with the 11nm chromatin fibre representing simple nucleosome arrays (so-called beads on a string)^6, 7^. A second model, originally proposed by Paulson and Laemmli^8, 9^ and supported by recent findings^10^, evokes a chromosome scaffold that comprises a continuous core of non-histone proteins running along the axis of the chromosome, from which radial loops of chromatin are attached. Although discrete in their architectures, both models envisage the formation of mitotic chromosomes as a multi-layered process requiring specialised proteins (such as condensin 1 and 2) and bespoke modifications to histone tails (such as histone H3S10 phosphorylation) that collectively drive spatial compression via close-range and more distant interactions^11^. While these interactions differ from those at interphase, it seems likely that at least some factors mediating local chromatin condensation during interphase might have similar or related roles in mitosis. The cohesin complex, for example, is required to keep newly replicated sister chromatids in close physical proximity during S/G2 to M phases of the cell cycle, but can also modulate interphase gene expression through local enhancer-promoter contact^12–17^.

Transmission of gene expression features from mother to daughter cells is also associated with DNA sequence-specific transcription factor binding through cell division^18^. Although it was originally thought that most sequence-specific transcription factors were likely to be displaced from chromosomes at mitosis^19^, subsequent studies focused on the Hsp70 gene promoter ^20^, or that have examined the dynamic distribution of GATA1, FOXAI and ESRRB in cycling cells^21–23^ have shown that these factors remain bound to mitotic chromosomes and occupy a subset of the genomic sites bound in interphase^23, 24^. In this setting, the continued binding of such factors has been proposed to ‘bookmark’ the mitotic genome, marking out specific genes for subsequent activity in daughter cells. Over the last few years, the number of mitotic bookmarking factors described has increased and now includes transcription factors associated with pluripotency (such as Oct4, Sox2 and Klf4) as well as more ubiquitously expressed chromatin-modifiers (such as Brd4, Mll, Ring1a and Bmi1)^23, 25–31^. Studies have also shown that the retention of DNA binding factors on mitotic chromosomes can occur through interactions with a variety of emergent features of condensed mitotic chromatin^32–35^, rather than exclusively through cognate DNA motifs. To comprehensively evaluate the repertoire and roles of mitotic chromosome-binding proteins in dividing pluripotent ESCs we wanted to move away from candidate-based analysis towards higher-throughput and unbiased approaches. Technical difficulties associated with successfully synchronising cells, ensuring that mitotic samples do not contain interphase contaminants, and a requirement to independently validate ‘chromosome-bound’ components from among heterogeneous mitotic lysates, have limited the performance of many conventional approaches. In addition, an increased appreciation that fixatives intended to stabilize or cross-link mitotic preparations can artificially displace factors from native mitotic chromosomes^28, 36, 37^, infer that prior studies might have substantially underestimated the repertoire of factors that bind to condensed mitotic chromosomes *in vivo*. To circumvent these issues, we directly isolated native (unfixed) mitotic chromosomes from dividing ES cells using Hoechst 33258 and Chromomycin A3 staining of DNA and flow cytometry to enumerate and sort specific chromosomes on the basis of their AT/GC content and forward scatter. This builds upon prior approaches^38–40^ enabling mitotic chromosome purification from different cell types and species. By performing LC-MS/MS analysis on conventional metaphase-arrested ESCs (lysates), in parallel with highly-enriched (sorted) chromosomes, we were able to catalogue the factors present in mitotic ESCs and discriminate chromosome-bound factors as being significantly enriched in chromosome-sorted fractions (Figure 1). From among 5,888 proteins reproducibly detected in mitotic ESC lysates, only around 10% (614) were significantly enriched on purified mitotic chromosomes. These included transcription factors (such as Sox2, Esrrb and Sall4), members of the structural maintenance of chromosomes family of proteins (such as Smc1), heterochromatin-associated proteins and repressive chromatin modifiers such as DNA methyl-transferases, PRC1 and PRC2. Importantly, we showed that individual mitotic chromosomes isolated from ESCs lacking PRC2 activity, DNA methylation or Mecp2, were significantly de-condensed relative to equivalent chromosomes from wild type (WT) ESCs, providing a functional validation for the roles of these components on mitotic chromosomes. In addition, metaphase chromosomes isolated from differentiated cells were more compact than equivalents isolated from ESCs and these isolated chromosomes remained responsive to biologically-relevant cues. *In situ* cleavage of endogenous cohesin complexes provoked de-condensation and a loss of structural integrity. Our study therefore illustrates a new approach in studying the properties of mitotic chromosomes and reveals an inventory of chromosome-bound proteins that comprise the mitotic signature of mouse pluripotent stem cells.

**Figure 1.**
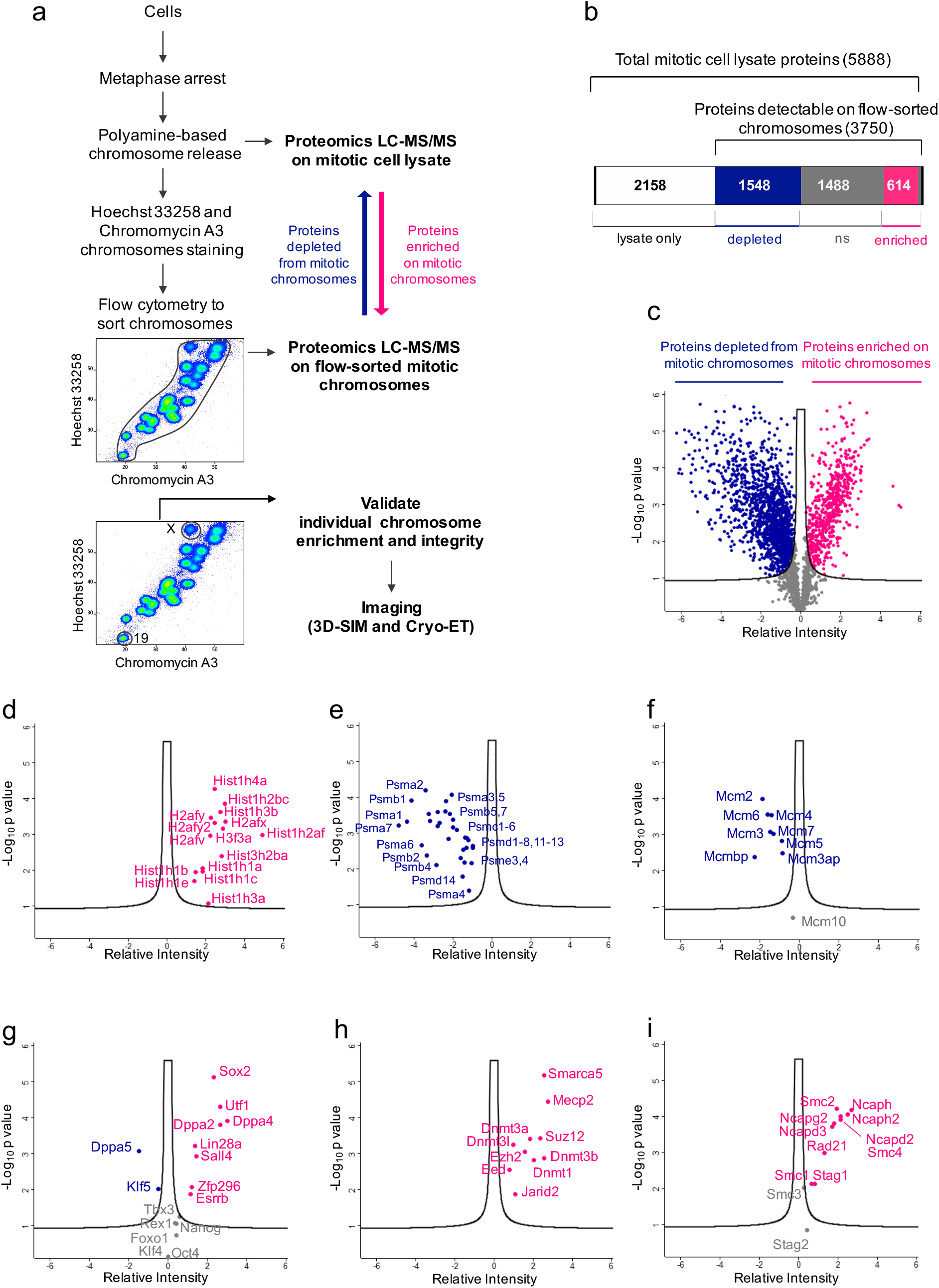
Proteins bound to ESC-derived metaphase chromosomes. (a) Scheme of experimental strategy used to isolate native metaphase chromosomes from ESCs and identify proteins bound to mitotic chromatin. Hoechst33258 and Chromomycin A3 bivariate Karyotype was assessed by flow cytometry and the gates used to sort all chromosomes, chromosomes 19, or the X chromosomes are indicated. Proteomic analysis was performed using LC-MS/MS on total mitotic cell lysate, or on flow-purified chromosomes, to identify proteins bound to native metaphase chromosomes. (b) Diagram of number of proteins identified by proteomic analysis as differentially detected in lysates and chromosome sorted samples. (c) Volcano plot of proteins detected as being significantly (FDR<0.01) enriched on sorted chromosomes relative to mitotic lysates. Proteins were plotted by relative intensity (LFQ intensity of Purified chromosomes – LFQ intensity of mitotic lysates) and significance (−Log p) using Perseus software. Volcano plots show (d) histones, (e) components of the proteasome, (f) DNA replication machinery, (g) pluripotency-associated transcription factors, (h) chromatin repressors or (i) SMC associate proteins, that are enriched (red), depleted (blue) or not significantly enriched (grey) on ESC mitotic chromosomes versus mitotic lysates.

## Results

### Isolation of native metaphase chromosomes from mouse ESCs

We adapted a protocol used previously to isolate unfixed mitotic chromosomes^40^. Briefly, rapidly dividing cultures of mouse ESCs were arrested in metaphase using demecolcine to achieve samples where most (85-90%) of cells were in M-phase as judged by PI labelling (Figures 1a and S1a). Condensed chromosomes were released using polyamine buffer, stained with Hoechst 33258 and Chromomycin A3 as described previously^40–42^, and examined by flow cytometry using a Becton Dickinson influx equipped with specialised air-cooled lasers (see detailed methods). This approach allows individual chromosomes to be discerned and either sorted en mass (upper plot, Figure 1a) or gated on individual chromosomes such as the X or chromosome 19 (highlighted separately in lower plot, Figure 1a). After sorting, the integrity of sorted chromosome was examined and verified by optical imaging using antibody to CENPA or using TRF1-YFP to confirm centromere and telomere number and location (Figure S1b).

### Analysis of proteins bound to isolated metaphase ESC chromosomes

To determine the proteins protein bound to native (unfixed) metaphase ESC chromosomes we performed a proteomic analysis using LC-MS/MS and analysed the data using the label-free quantification (LFQ) algorithm within the MaxQuant software platform. In these experiments we compared samples with equivalent numbers of ESC metaphase chromosomes (10^7^) before and after chromosome sorting, in three biological replicates. This led to the identification of 5888 proteins in mitotic lysates of which 5436 were identified with two or more razor or unique peptides per protein at a 1% false discovery rate (FDR). Our rationale was that chromosome-bound factors should be enriched in sorted samples, while factors that were not chromosome-associated would be depleted. In the sorted fractions we detected 3750 proteins (Figure 1b) that were either significantly enriched as compared to mitotic lysates (614, red), were depleted relative to mitotic lysates (1548, blue) or showed no statistical difference between the two (1488, grey) (Figures 1b and 1c). Consistent with our expectations, we found histones significantly enriched in the sorted samples (Figure 1d, Red) whereas components of the proteasome and members of the minichromosome maintenance complex (Mcm) were depleted from mitotic chromosomes (Figures 1e and 1f respectively, blue). Mcm components are critical for initiating DNA synthesis but are known to dissociate from chromosomes once DNA synthesis has occurred in S-phase and cells transit into G2^43^. The relative depletion of Mcm factors among mitotic chromosome-sorted samples provides further confidence that chromosome-bound and non-bound proteins can be discriminated using this approach. Further analysis of the proteomic data (Figure S1c) showed that chromatin remodelling complexes, chromosome architectural proteins and selected DNA binding proteins were enriched in chromosome-sorted samples. Importantly, among the transcription factors that regulate ESC pluripotency and differentiation, enrichment on mitotic chromosomes was selective (Figure 1g). For example, Utf1, a transcription factor required for the proper differentiation of embryonic carcinoma and embryonic stem cells^44^ was enriched in sorted mitotic chromosome samples, while Nanog, Oct4 (Pou5f1) and Klf4, although detected, showed no significant enrichment on sorted chromosomes. Esrrb and Sox2, two previously characterised bookmarking factors^23, 24, 26–28^ were significantly enriched in chromosome-sorted samples, while factors such as Dppa5 and Klf5 were significantly depleted from chromosome-sorted fractions, implying that they might be evicted from chromosomes during mitosis. Antibody-mediated labelling confirmed abundant Sox2 bound to isolated mitotic ESC chromosomes (Figure S1d), while Oct4 and Nanog appeared low or undetectable, consistent with the proteomic analysis. Proteins associated with gene repression and heterochromatin formation including the HMTases Suv39h1 and Suv39h2 and components of PRC1 and PRC2 such as Jarid2, Eed and Ezh2, were all significantly enriched in chromosome-sorted samples (Figure 1h and supplemental table1). The DNA methyltransferases Dnmt1, Dnmt3a and Dnmt3b also showed co-enrichment after chromosome sorting, suggesting that most of the chromatin machinery required to sustain H3K9me3, H3K27me3 and 5mC marking of the genome remains bound to ESC chromosomes through mitosis. Likewise, the methyl-CpG-binding protein Mecp2, and the SWI/SNF related protein Smarca5 were also significantly enriched in chromosome-sorted samples (Figure 1h). Smarca5 is an important chromatin remodelling factor that is involved in establishing regularly spaced nucleosomes, critical for DNA replication and repair, and associated with both positive and negative transcriptional outcomes. Smarca5 has been shown to interact with Rad21^45^, and enrichment of Rad21, Smc1, Smc2 and other SMC-associated proteins was evident among chromosome-sorted samples (Figure 1i, red), consistent with these factors also remaining bound to ESC metaphase chromosomes. Rad21 binding to isolated ESC chromosomes was confirmed by antibody labelling (Figure S1d) showing a centromeric distribution with lower abundance along chromosome arms.

### DNA methylation and PRC2 activity keep mitotic chromosomes compact

To investigate the relevance of these chromatin-based repressors we examined mitotic chromosomes from ESCs that lack DNA methylation^46^ or PRC2 activity^47^. Although the overall distribution of mitotic chromosomes isolated from WT and mutant ESCs (*Dnmt1,3a,3b*^−/−^ or *Eed*^−/−^) was similar (Figure 2a), a close inspection revealed differences in the size and shape of these mitotic chromosomes. To accurately measure this, we separately purified two representative mitotic chromosomes (19 and X) from WT and mutant ESCs, using Hoechst 33258 and Chromomycin A3 staining and flow sorting as described previously (Figure 1a). DNA-FISH with mouse chromosome 19- or X-specific paints (Figure S2a) confirmed 99-100% sample purity. The size of individual chromosomes was estimated by microscopy using standard imaging software to determine the total chromosome area and estimate the size of DAPI-bright pericentric domains (Figure S2b). These analyses indicated that mitotic 19 and X chromosomes isolated from *Dnmt1,3a,3b*^−/−^ (23.4±4.1, 43.6±6.9μm^2^ respectively) or *Eed*^−/−^ (24.4±4.8, 42.6±4.7μm^2^ respectively) ESCs were significant larger than equivalent chromosomes isolated from WT ESCs (18.4±3.1, 34.6±5.3μm^2^ respectively) (Figures 2b and 2c). In contrast, mitotic chromosomes isolated from a Sox2-deficent ESC line^48^ were comparable in size and shape to equivalent mitotic chromosomes isolated from WT ESCs (Figure 2b and 2c). These data suggest that DNA methylation and PRC2 activity are important for efficient chromosome condensation in mitosis. Mitotic chromosomes from WT ESCs showed a distribution of H3K27me3 along the entire chromosome, with centromeres decorated by histone H3K9me3 (Figure S2e). *Dnmt1,3a,3b*^−/−^ mutants lacked 5mC and had reduced levels of both H3K9me3 and H3K27me3 (mid panel Figure S2c-d and middle panel S2e), while *Eed* mutant lacked H3K27me3 but retained H3K9me3 and 5mC (Figure S2c-d and right panel S2e).To exclude that chromosome de-compaction seen in PRC2-deficient ESCs was not due to inadvertent secondary effects, we examined *Eed*^−/−^ ESCs transfected with a BAC that contains Eed (clone B1.3BAC) that had previously been shown to restore PRC2 activity and H3K27me3^47^. As shown in Figure 2d, metaphase chromosomes isolated from Eed-rescued (*Eed-BAC*) were similar in size and shape to equivalent chromosomes from WT ESCs, confirming that the de-compaction seen in Eed-null chromosomes was fully reversed by restoring PRC2 activity.

**Figure 2.**
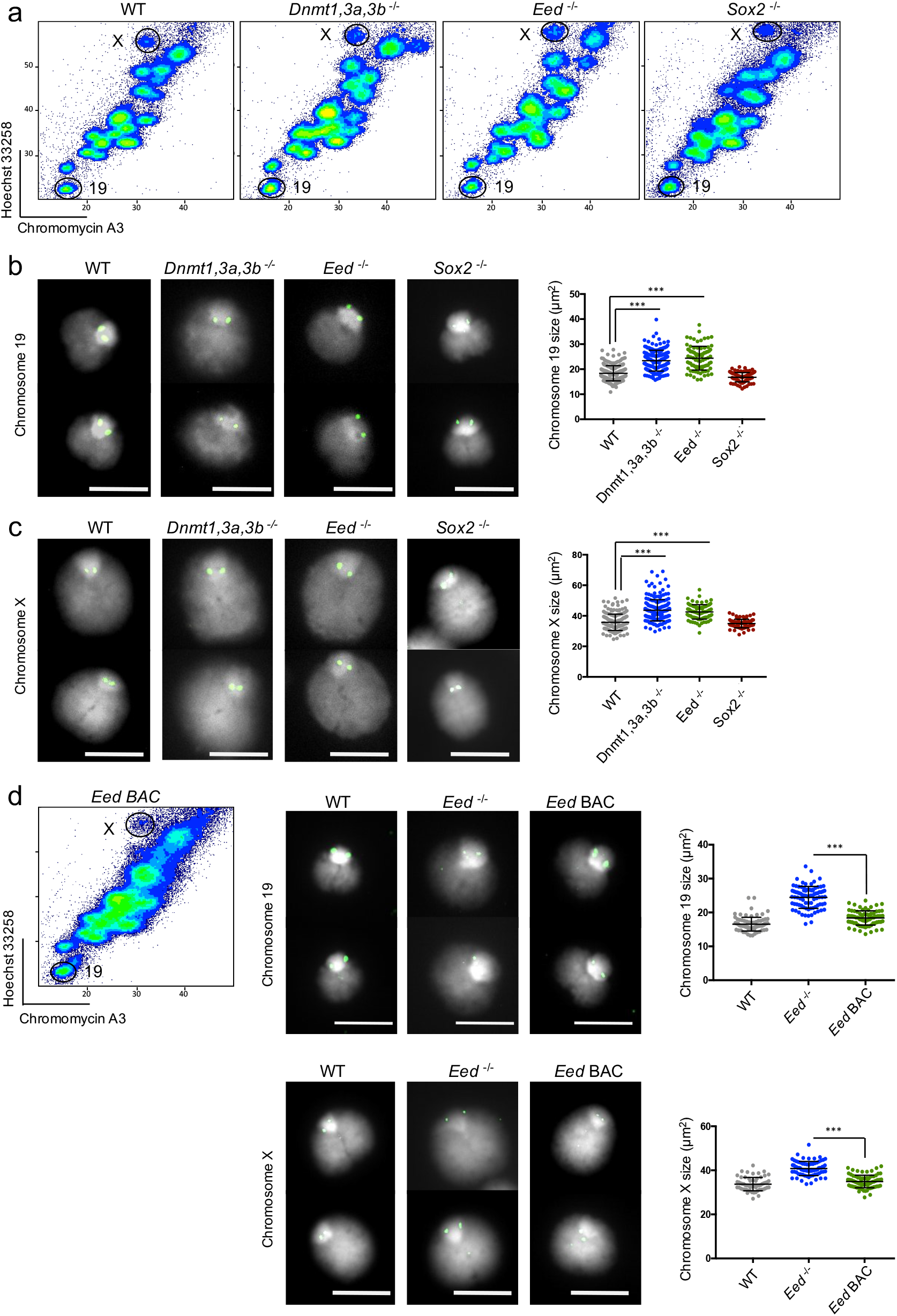
Increased size of ESC mitotic chromosomes that lack DNA methylation or PRC2 activity. (a) Flow karyotype of mitotic chromosomes isolated from WT ESCs or mutant ESCs that lack Dnmt1,3a,3b, Eed or Sox2. Gates used to isolate chromosome 19 or X are indicated. Representative images of mitotic chromosomes 19 (b) and X (c) from different ESCs are shown, where DAPI stain (light grey) and Cenpa label (green) indicate the chromosome body and centromere respectively, scale bar =5μm. Chromosome size was calculated by measuring a minimum of 100 chromosomes for each ESC line, mean ± SD are indicated. (d) Flow karyotype and mitotic chromosome size of *Eed*^−/−^ ESCs before, and after restoring *Eed* expression (*Eed BAC*). Representative images show chromosome 19 (top panel) and X chromosome (lower panel) isolated from WT, Eed-deficient (*Eed*^−/−^) and rescued (*Eed BAC)* ESCs, where chromosome size was measured for a minimum of 100 chromosomes for each cell line, and mean± SD values are shown. Scale bar=5μm. Asterisks show statistical significance estimated by unpaired two-tailed Student’s t test ***<0.001.

### Mecp2 contributes to the mitotic compaction of autosomes in ESCs

To determine whether the methyl binding protein Mecp2 contributes to mitotic chromosome condensation we examine ESCs lacking Mecp2^49^. Mecp2 has been implicated in regulating chromatin architecture at a range of different levels, from the juxtaposition of nucleosome arrays, to condensing pericentric heterochromatin, and is reported to interact with a plethora of partners independent of binding methylated CpG (reviewed in^50^). Comparing metaphase chromosomes 19, 3 and X isolated from parental male Mecp2-expressing (*Mecp2*^*lox/y*^) and Mecp2-deficient (*Mecp2*^−/*y*^) ESCs we observed a significant increase in the sizes of both autosomes in the absence of Mecp2, but no apparent change in the size of active X chromosomes (Figure 3a and 3b). Although CpG methylation is found on active and inactive X chromosomes, published ChIPseq analysis indicates that MeCp2 is less abundant on X and Y chromosomes than on autosomes (Figure S3a and S3b) in ESCs and in neurons^51, 52^. Depletion of Mecp2 provoked a profound change in the distribution of histone H3K9me3 along metaphase chromosomes. In particular, dense H3K9me3 labelling that normally is confined to pericentric (DAPI-intense) regions on metaphase chromosomes, extended along the arms of autosomes in the absence of Mecp2 (Figure 3c, lower panel, quantified in S3c and S3d). This spread in H3K9me3 distribution beyond DAPI-intense regions, was not evident in equivalent chromosomes isolated from *Dnmt1,3a,3b*^−/−^ or *Eed*^−/−^ ESCs, and H3K9me3 was much reduced in *Dnmt1,3a,3b*^−/−^ ESCs (Figure S2e, green).

**Figure 3.**
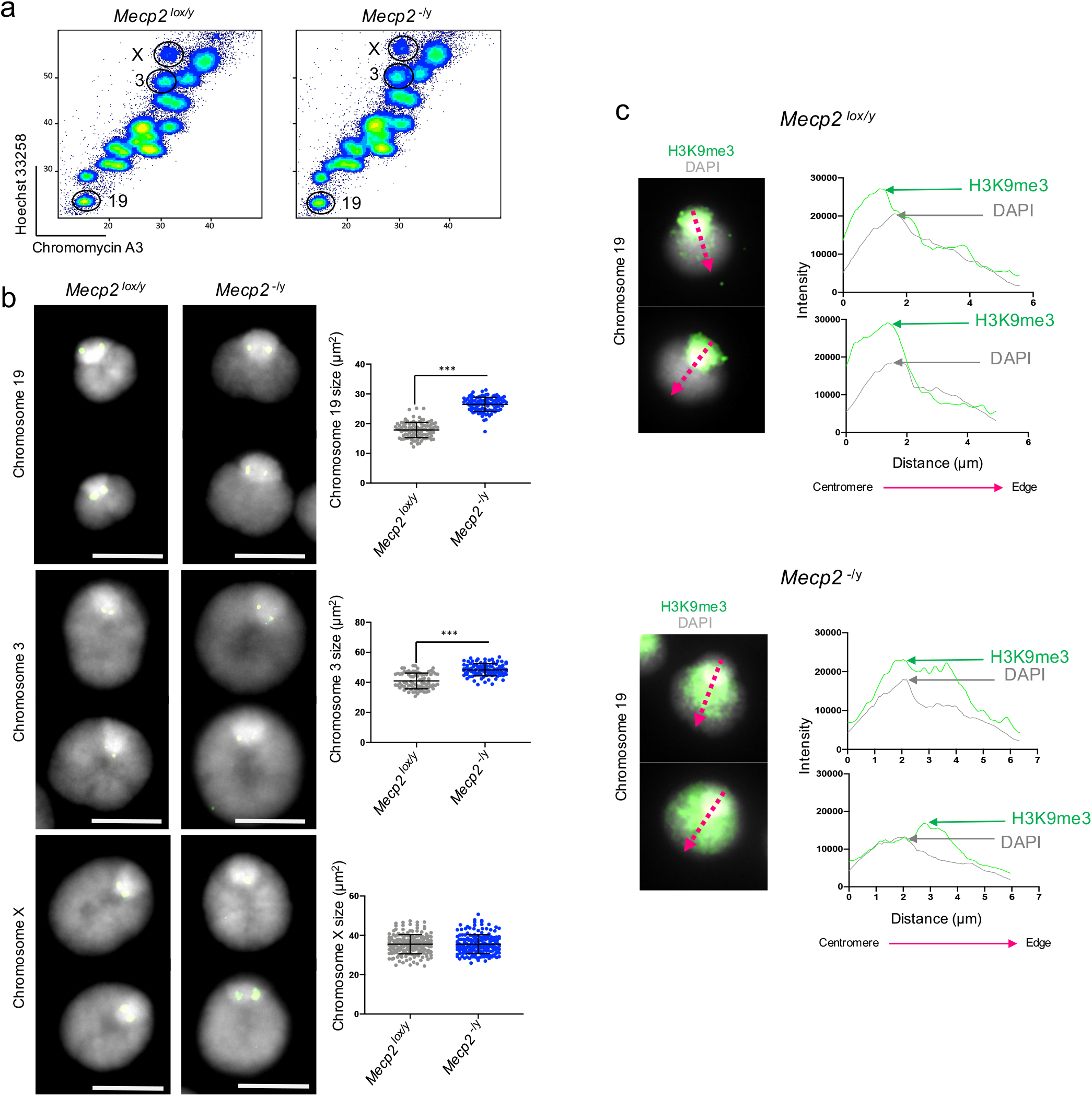
Increased size of mitotic chromosomes in ESCs lacking Mecp2. (a) Flow karyotype of mitotic chromosomes isolated from *Mecp2^lox/y^* or *Mecp2*^−/*y*^ ESCs. Gates used to isolate chromosome 19, 3 or X are indicated. Representative images of mitotic chromosomes 19, 3, and X (b) from ESCs *Mecp2*^*lox/y*^ and *Mecp2*^−/*y*^ are shown, where DAPI stain (light grey) and Cenpa label (green) indicate the chromosome body and centromere respectively, scale bar =5μm. Chromosome size was calculated by measuring a minimum of 100 chromosomes for each ESC line, mean ± SD are indicated. Asterisks indicate statistical significance estimated by unpaired two-tailed Student’s t test ***<0.001. (c) Representative images of immunofluorescence labelling of histone H3K9me3 (green) and DAPI (light grey) on mitotic chromosome 19 from *Mecp2*^*lox/y*^ and *Mecp2*^−/*y*^ ESCs. Distance distribution plots of H3K9me3 (green) and DAPI (grey) intensities were measured for each chromosome along a central selected axis (red) from the centromere to the distal edge of the chromosome arm.

### Mitotic chromosome size depends upon cell context and differentiation state

As PRC2 activity and DNA methylation are required for successful differentiation but not for ESC self-renewal or pluripotency (reviewed in^53, 54^), we asked whether mitotic chromosomes isolated from differentiated cells and ESCs were similar. To examine this, we isolated individual chromosomes from metaphase arrested mouse ESCs (Figure 4a), pre-B cells (Figure 4b), mouse cardiomyocyte HL-1 cells (Figure 4c) and primary embryo fibroblasts (Figure 4d). Although the success of metaphase arrest varied between the different cell types (45%-90%), by applying a FACS-based approach we were able to isolate and purify mitotic chromosomes, irrespective of differences arising from cell cycle synchronisation or karyotype complexity. Using chromosomes 19 and X as representatives, we found that mitotic chromosomes isolated from differentiated pre-B cells, cardiomyocytes and fibroblasts, were significantly smaller than equivalents isolated from ESCs (Figure 4e and 4f). As these size differences could reflect constraints imposed earlier in the cell cycle, for example by the size of nuclei, we measured the diameter of G1- and G2-phase nuclei in each of the different cell types (Figure S4a). This comparison showed similar nuclear sizes in each of the different cell types examined in G1 (Figure S4b), or in G2-phases (Figure S4c).

**Figure 4.**
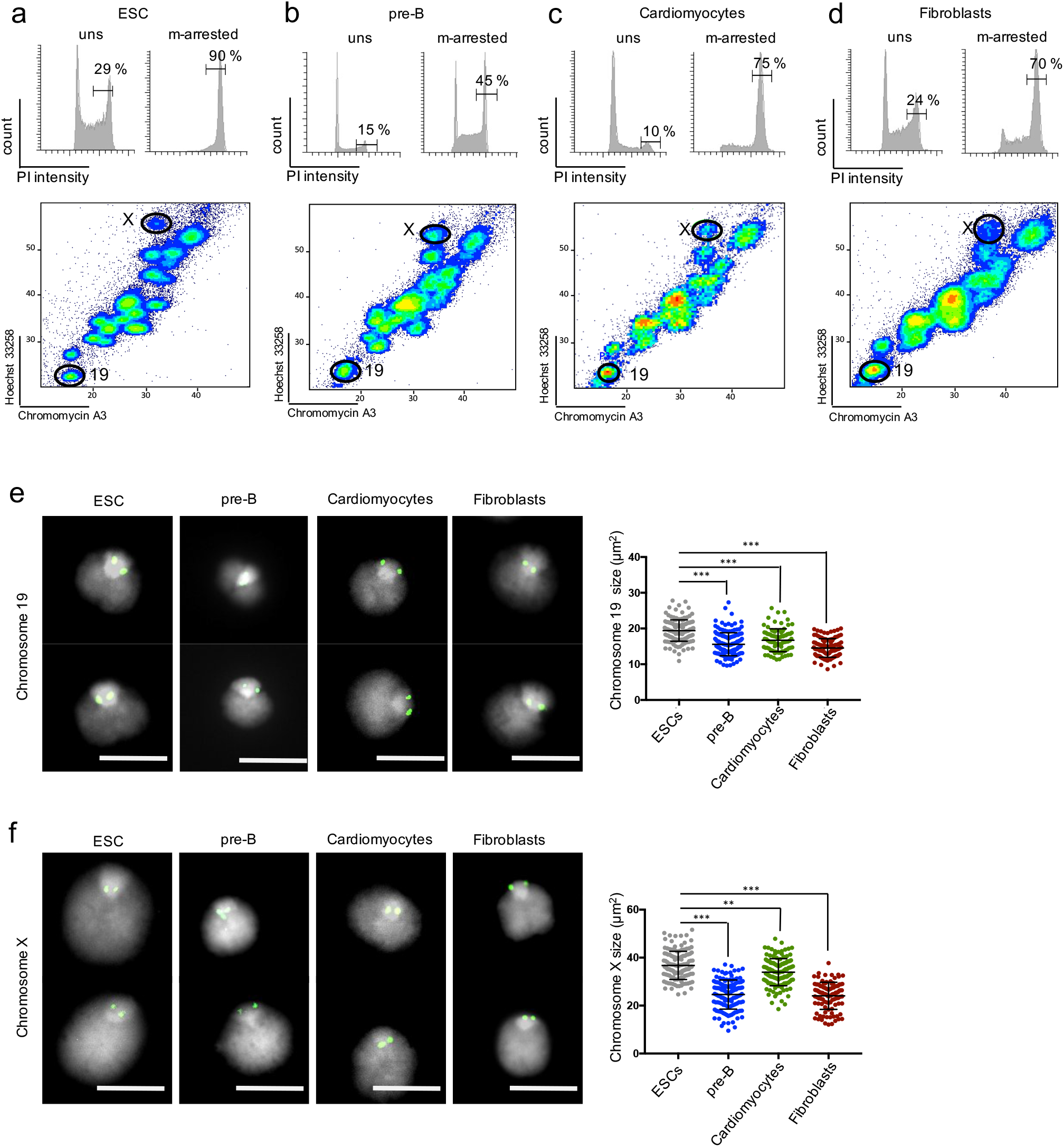
Native mitotic ESC chromosomes are larger than equivalents isolated from differentiated cells. Cell cycle profiles of mouse ESCs (a), pre-B cells (b), cardiomyocytes (c) and embryonic fibroblasts (d), were determined by staining with propidium iodide (PI). Left panel shows unsynchronized (uns) cells, right panel shows samples 6-12 hours after treatment with demecolcine (m-arrested), where values indicate percentage cells in G2/M cells. Lower panel shows flow karyotype of demecolcine-treated cells and the gates used to isolate chromosomes 19 and X. Representative images of native mitotic chromosomes 19 (e) and X (f) isolated from mouse ESCs, pre-B cells, cardiomyocytes (HL-1) and embryonic fibroblasts. DAPI stain (light grey) and Cenpa (green) labelling are shown, scale bars=5 μm. Chromosome size was determined by measuring chromosome area for a minimum of 100 chromosomes for each cell type, and calculating mean ± SD. Asterisks indicate statistical significance estimated by unpaired two-tailed Student’s t test ***<0.001.

### Isolated metaphase chromosomes are sensitive to *in situ* cohesin cleavage

A major potential advantage of purifying native chromosomes from cells in mitosis is that these unfixed chromosomes may respond to appropriate biological cues when applied ‘*in situ*’. To test this, we asked whether experimental cleavage of cohesin could significantly alter the structure of isolated chromosomes. Cohesin complexes are composed of Smc1, Smc3, the kleisin Scc1, and one of three auxillary subunits^13^ and are required to keep sister chromatids together from DNA replication until mitosis, as well having roles in interphase genome organisation, gene transcription and DNA repair^12–17^. Proteomic analysis (Figure 1i and S1c) indicate that components of the cohesin complex bind mitotic chromosomes in different cell types^25, 55, 56^ and are dynamically regulated so that the majority of cohesin dissociates from chromosome arms during prophase. Some cohesin remains bound at centromeres until anaphase when separase cleaves the kleisin subunit^13, 57, 58^. We used a pre-B cell line that expresses a cleavable form of Rad21 (Rad21-TEV-myc) to investigate the impact of cohesin complex dissolution^59, 60^. We first examined the distribution of Rad21 on native mitotic chromosomes isolated from pre-B cells and from ESCs and, as anticipated, detected substantial Rad21 binding at centromeric regions (as illustrated for chromosome 19, Figure 5a and S5a). We then performed Hi-C analysis of flow-sorted mitotic chromosomes from pre-B cells to confirm that the 3D chromosome contacts present at interphase (Figure S5b, upper panels) were lost from mitotic chromosome samples (Figure S5b, lower panels), consistent with previous reports^10, 61^. We then isolated native mitotic chromosomes from WT and *Rad21*^*Tev/Tev*^ pre-B cell lines (Figure S5c) and examined the impact of cohesin cleavage induced by TEV protease (Figures 5b). Briefly, metaphase-arrested WT or *Rad21*^*Tev/Tev*^ pre-B cells were used to sort mitotic chromosomes *en mass*, or gated for chromosome 19, and these samples were treated with TEV enzyme (0.06U/ml for 4 hours) or with buffer alone and imaged using advanced optical microscopy or cryo-electron tomography (cryo-ET) (Figure 5c shows the experimental design). TEV treatment resulted in efficient and selective cleavage of Rad21-Tev, as verified by Myc immunofluorescence labelling (Figure 5d) and Western blotting (Figure S5d). TEV-induced cohesin cleavage also resulted in a significant increase in the size of mitotic chromosome 19, compared with either untreated (-TEV) or TEV-treated chromosomes derived from WT pre-B cells (Figure 5e). Increased mitotic chromosome 19 size was accompanied by a de-condensation at DAPI-bright pericentric domains (arrowed). Further cryo-ET analysis of two independent experiments (Figure 5f) confirmed that TEV-induced cohesin cleavage resulted an increase in the size of mitotic (*Rad21*^*Tev/Tev*^) chromosome 19, and a widespread de-condensation was evident in representative 3D images (supplemental videos 1 and 2).

**Figure 5.**
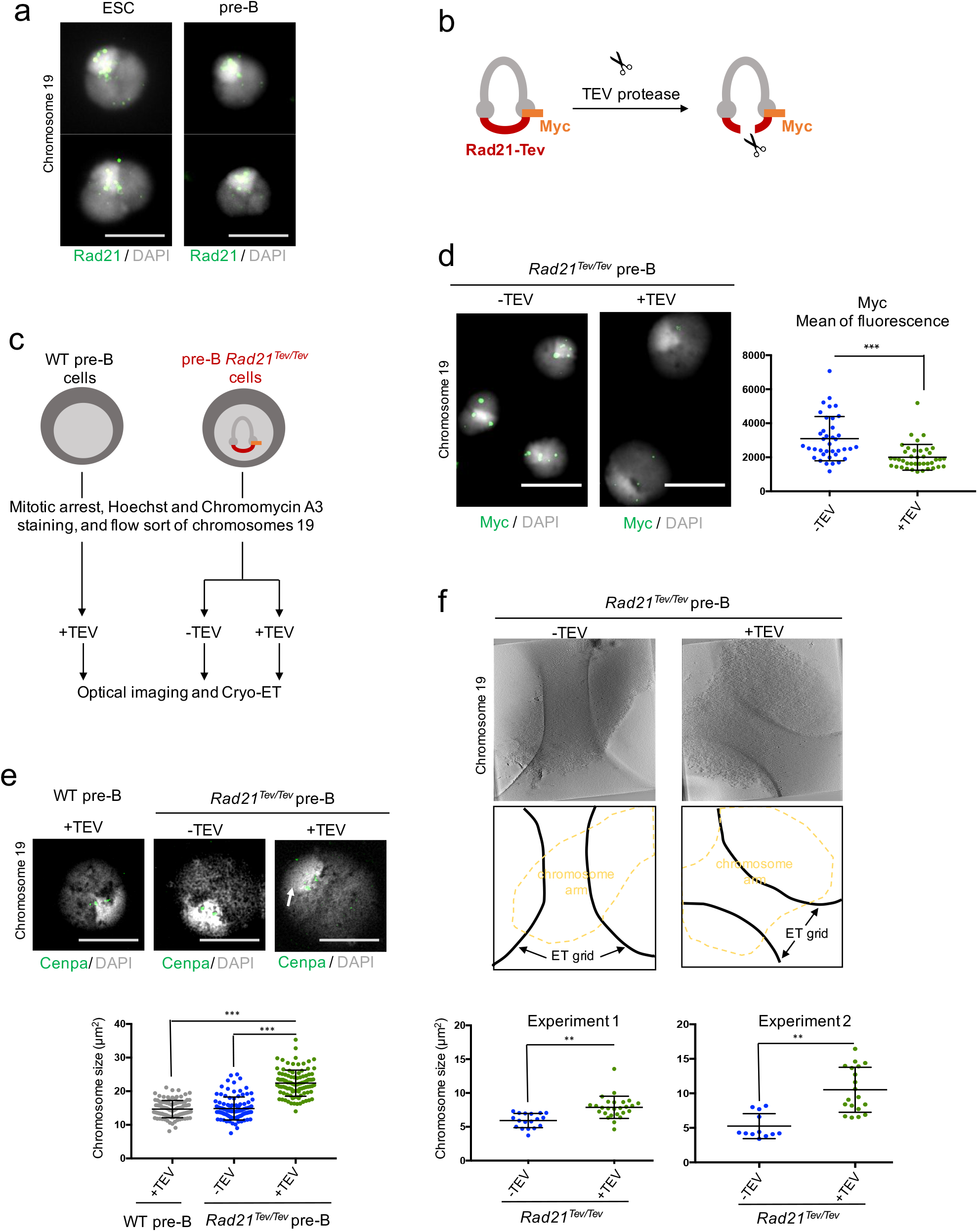
Experimentally induced cleavage of cohesin alters mitotic chromosome size. (a) Rad21 labelling (green) shows cohesin bound to native mitotic chromosome 19 isolated from ESCs or pre-B cells, scale bars=5μm. (b) Experimental strategy used to cleave cohesin using TEV protease; illustrated is a cohesin ring containing Smc1, 3, and Rad21-Tev-Myc. (c) Scheme used to isolate and image mitotic chromosomes from WT and Rad21-Tev-myc pre-B (*Rad21*^*Tev/Tev*^) cells. Mitotic chromosomes of WT pre-B cells or from *Rad21*^*Tev/Tev*^ pre-B cells were purified by flow cytometry and incubated with or without TEV protease. (d) Myc labeling (green) of *Rad21*^*Tev/Tev*^ purified chromosome 19 shows reduced Myc levels after treatment with TEV protease (images left, and quantified by intensity, right), scale bars=5μm, where at least 30 chromosomes were examined for each condition and mean intensity values ± SD are shown. (e) Representative super-resolution SIM images of purified mitotic chromosome 19 isolated from WT or *Rad21*^*Tev/Tev*^ pre-B cells, treated with TEV *in situ* (+TEV) or with buffer alone (-TEV). Chromosome size was measured for a minimum of 100 chromosomes for each condition, values indicate mean ± SD, scale bars=2.86μm. (f) Representative slices through cryo-electron tomograms (Cryo-ET) of chromosome 19 isolated from *Rad21^Tev/Tev^* pre-B cells and treated with TEV *in situ* (+TEV) or with buffer alone (−TEV) (top panel) and Cryo-ET image explanation (middle panel). Graphs show chromosome size, calculated as area measurements from 2D electron microscopy images. Values from two independent experiments are shown. Asterisks indicate statistical significance estimated by unpaired two-tailed Student’s t test. ***<0.001, **<0.01.

## Discussion

The isolation and purification of mitotic chromosomes by flow cytometry offers a new approach for studying chromosome structure and dissecting the interplay between transcription factors and chromatin that convey cellular memory through mitosis. We show that the size of individual metaphase chromosomes differs between different cell types, something that we believe has not been reported previously. Native mitotic chromosomes purified from ESCs were much less condensed than equivalents isolated from either lymphocytes, cardiomyocytes or fibroblasts. The idea that chromosomes of pluripotent ESCs might be more ‘loosely-packed’ than somatic equivalents is consistent with previous studies in interphase in which electron spectroscopic imaging (ESI) revealed that ESCs and cells of the mouse early-epiblast (E3.5) lack the compact chromatin domains that characterise differentiated lymphocytes, liver and kidney cells^62^. ESI studies have also shown that at later stages of development (E5.5) epiblast cells loose the dispersed 10nm chromatin fibres that are so-called architectural hallmarks of pluripotency^63^. Our ability to purify and compare mitotic chromosomes from stem and differentiating cells will be important for future studies aimed at understanding the basis of these dynamic changes in chromosome structure, and mapping changes in mitotic bookmarking at key developmental transitions. We have also shown that native metaphase chromosomes purified by FACS, remain responsive to appropriate biological cues. As an example, we show that *in situ* cleavage of residual cohesin that is bound to centromeric domains on metaphase chromosomes, provokes de-condensation and the loss of structural integrity. This infers that metaphase chromosomes isolated in their native state remain sensitive to regulators of chromosome architecture, a finding that should enable the mechanisms of chromosome condensation and de-condensation to be more closely observed. We examined the repertoire of proteins bound to metaphase chromosomes in pluripotent ESCs, using FACS and a quantitative LC-MS/MS approach to discriminate proteins that were significantly enriched on sorted chromosomes, versus ‘unbound’ proteins in mitotic lysates. This revealed a repertoire of >600 proteins that remain bound to metaphase ESC chromosomes, many of which have established roles in chromatin organisation, DNA and nucleosome packaging, cell cycle and chromosome architecture and function. Three subsets of transcription factors relevant for pluripotency were discerned in mitotic ESCs that showed either a significant enrichment on metaphase chromosomes versus lysates (eg. Sox2, Utrf1, Dppa4, Dppa2, Sal4, Esrrb), were depleted (eg. Dppa5, Klf5) or were similarly represented in both (eg. Oct4, Nanog, Klf4). The first group includes Esrrb and Sox2, transcription factors that were previously been shown to remain bound throughout mitosis in ESCs and implicated in mitotic bookmarking^23, 27, 28^. The later two subgroups included candidates evicted from condensing metaphase chromosomes or that were in dynamic flux so that they appeared to be equivalently distributed between mitotic lysate and chromosomes samples. Examples of this include Oct4, Nanog and Klf4 that were detected but showed no significant enrichment after chromosome sorting. Studies by others have shown that Oct4, Nanog and Klf4 can remain bound to metaphase ESC chromosomes^26^, albeit at low levels (Figure S1d). Conceivably differences in these results could reflect subtle changes in transcription factor concentration and DNA binding (on rates) at isolated mitotic chromosomes that may impact occupancy^64^.

Proteomic analysis also showed that many of the core components of Polycomb Repressor Complex 1 (Rnf2, Pcgf6, Cbx2, Phc1) and Polycomb Repressor Complex 2 (Eed, Ezh2, Suz12) that are responsible for catalysing histone H2AK119 mono-ubiquitination and histone K3K27 tri-methylation respectively, were enriched on metaphase ESC chromosomes. Likewise, DNA methyl-transferases (Dnmt1, Dnmt3a and Dnmt3b) and the methyl-CpG-binding protein Mecp2 were also significantly enriched on metaphase chromosomes as compared to lysates. Previous studies had suggested that Polycomb group proteins PC, PH, PSC and BMI1 dissociate from condensing chromosomes from prophase to metaphase in *Drosophila* embryos and in human primary cells^65, 66^. Studies of living, rather than fixed samples, indicated that although GFP-tagged PC fusion proteins are depleted from mitotic chromosomes relative to interphase, some PC complexes remained bound^67^. Our results were obtained using unfixed but highly purified metaphase ESC chromosomes and resemble data from a recent study of chromatin-bound changes through the cell cycle in human glioblastoma T98G cells^25^ showing that chromatin repressor complexes remained bound throughout mitosis. Among chromatin activator complexes, including those previously linked to stem cell self-renewal and pluripotency^68–70^, we found a wide range of distributions. Some Trithorax (Trx) family members appeared to be enriched on metaphase chromosomes (Kmt2a, Kmt2b, similar to Chd1, Chd2, Chd4, Chd8) while others appeared more abundant in mitotic lysates (Brd4, Ash2l, Wrd5, Wrd82). Previous studies of Trithorax-family proteins during mitosis have reported that MLL can remain bound to^31^ or dissociate from mitotic Hela chromosomes, being rebound upon exit from mitosis^71^. Studies of chromatin activator complexes in *Drosophila* have also suggested that trithorax proteins such as Ash1, can remain bound to mitotic chromosomes in the embryonic blastoderm while Polycomb-group proteins such as EZ, PHO and PC dissociate^72^. While it is possible that complexity and functional redundancy within the trithorax family members^73^ may explain the often discrepant reports of occupancy during mitosis, our data underscore the importance of both cellular context and native (unfixed) samples in defining the metaphase chromatin repertoire.

We demonstrate that Dnmt1,3a,3b, Mecp2, PRC1 and PRC2 repressors remain bound to metaphase chromosomes in pluripotent ESCs. To assess their functional relevance, we examined chromosomes from ESCs lacking DNA methylation, Mecp2, or PRC2 activity (and associated H3K27me3), as compared to those lacking the transcription factor Sox2. Genome-wide loss of DNA methylation or H3K27me3 resulted in mitotic chromosomes that were larger and less compact than WT equivalents. At first sight this is surprising since in interphase reduced DNA methylation is reported to affect nuclear organisation, histone modifications and linker histone binding, but does not directly alter chromatin compaction^74^. Furthermore, at the nucleosome level, *in vitro* studies have indicated that DNA methylation alone does not induce chromatin compaction^75, 76^. This suggests that the de-compaction of metaphase chromosomes seen in *Dnmt1, 3a, 3b*-null ESCs most likely stems from secondary changes that serve to relax the higher order structure of chromatin^6^, perhaps through altered histone H1 binding or an impaired recruitment of Mecp2. Mecp2 is a multifunctional protein that can bind both methylated and un-methylated DNA, can compete with histone H1, and has been shown to directly mediate nucleosome oligomerisation and compaction *in vitro*^*77*^. Consistent with this, metaphase chromosomes isolated from *Mecp2*-null ESCs also showed a de-condensation of autosomes relative to WT equivalents. The size of the X chromosome, that is active in WT male ESCs and relatively depleted of Mecp2 as compared to autosomes, was however unaffected. In post mitotic neurons Mecp2 depletion is reported to provoke an increase, rather than a decrease in chromatin compaction, associated with histone H4K20me3 being repositioned to pericentric heterochromatin territories previously occupied by Mecp2^78^. Our experiments have exclusively examined chromosomes during mitosis, where prior withdrawal of DNA methylation, PRC2-mediated H3K27me3, or Mecp2 each separately induced an increase in chromosome size, consistent these heterochromatin-associated features being important for keeping chromosomes compact. Taken together these experiments not only provide a functional validation for chromatin repressors being bound at metaphase chromosomes in ESCs but also highlight the importance of an interplay between chromatin remodellers and trans-acting factors for conveying mitotic memory.

## Materials and Methods

### Cells

ESCs used in this study were wild type (WT) E14Tg2a^48^, Sox2-deficient (clone 2O5)^48^, *Dnmt1,3a,3b*^−/−*46*^, *Eed*^−/−^ (clone B1.3), rescued Eed null (Eed BAC)^47^, and floxed Mecp2 ESC clones (*Mecp2*^*lox/y*^ and *Mecp2*^−/*y*^). These clones were generated by transfecting a floxed Mecp2 line with Cre. *Mecp2*^−/*y*^ clone has expressed Cre and deleted the floxed segment, *Mecp2^lox/y^* clone has been unaltered. ESCs were cultured on 0.1% gelatine-coated plates. Cells were grown in KO-DMEM medium supplemented with 15% FCS, non-essential amino acids, L-glutamine, 2-mercaptoethanol, antibiotics and 1000 U/ml of leukaemia inhibitory factor. Abelson-transformed pre-B cell lines (WT and *Rad21*^*Tev/Tev*^ pre-B cells were cultured in IMDM medium supplemented with 10% FCS, 2mM L-glutamine, antibiotics, and 50 μM 2-mercaptoethanol, and non-essential amino acids. These cell were derived in our lab from transgenic Rad21-Tev-Myc mice^60^. These mice express a modified and functional Rad21 protein containing three Tev cleavage sites within the flexible polypeptide connecting the N-terminal and C-terminal domains. A Myc tag was added together with the Tev sites and allow us to follow the cleavage and the chromosomal localisation of Rad21 by an anti-Myc antibody. Transgenic mice were obtained from Nasmyth laboratory (University of Oxford, Department of Biochemistry). Mouse cardiomyocyte cells (HL-1) were cultured in Claycomb medium (Sigma-Aldrich) supplemented with 10% of FBS (F2442, Sigma-Aldrich), 10 μg/ml Penicillin and Streptomycin, 2mM L-glutamine, and 0.1mM Norepinephrine. Mouse embryonic fibroblasts (MEFs) were cultured in DMEM supplemented with 10% of FCS, 10 μg/ml Penicillin and Streptomycin, 2mM L-glutamine.

### Mitotic arrest and cell cycle analysis

24h after passaging the cells, demecolcine (D1925, Sigma-Aldrich) was added to the culture media to a final concentration of 0.1μg/ml. ESCs and pre-B cells were incubated with demecolcine for 6h. Fibroblast and cardiomyocyte cells were incubated for 12h at 37°C. Cells were collected before and after metaphase arrest (after mitotic shake off). 10^6^ cells were fixed with ice-cold 70% ethanol, overnight at −20°C. Prior to staining, cells were washed twice with PBS and resuspended in the staining buffer containing 0.05 mg/ml of Propidium Iodide (PI), 1mg/ml RNaseA, and 0.05% of NP40. Samples were incubated 10min at RT and 20min on ice. PI signal was analyzed in a linear mode using BD LSRII flow cytometer and BD DIVA software.

### Chromosome preparation and flow sorting

Chromosomes were prepared using a polyamine-based method as described previously^40, 41^. Mitotic cells were collected by mitotic shake off. The cells were centrifuged at 1200rpm for 5min at room temperature (RT). The cell pellets were gently resuspended in 5-10ml of hypotonic solution (75 mM KCl, 10mM MgSO4, 0.5mM Spermidine, 0.2mM Spermine; pH=8) for 15min at RT. The swollen cells were then centrifuged at 1500rpm for 5min at RT and resuspended in 1-3ml of freshly prepared ice-cold polyamine isolation buffer (15 mM Tris-HCl, 2 mM EDTA, 0.5 mM EGTA, 80 mM KCl, 3mM DTT, 0.25% Triton X-100, 0.2 mM spermine and 0.5 mM spermidine; pH= 7.5). After 15 min of incubation on ice, the chromosomes were released by vortexing vigorously for 20-30s. To increase chromosome recovery, the suspension was passed through a 22-gauge needle using a 1 ml syringe. Chromosomes suspension were centrifuged for 2min at 1000rpm, RT. Then, the supernatant containing mitotic chromosomes was filtered using a 20-μm mesh filter into a 15-ml Falcon tube. Chromosomes were stained at 4°C overnight with 5μg/ml of Hoechst 33258, 50μg/ml of Chromomycin A3 and 10mM MgSO4. At least 1h prior chromosome sorting, sodium citrate and sodium sulfite were added to chromosome suspensions to a final concentration of 10mM and 25mM respectively. Chromosomes were examined by flow cytometry using a Becton Dickinson Influx equipped with spatially separated air cooled lasers. Hoechst33258 was excited using a (Spectra Physics Vanguard) 355nm laser with a power output of 350 mW. Hoechst33258 fluorescence was collected using a 400nm long pass filter in combination with a 500nm short pass filter. Chromomycin A3 was excited using a (Melles Griot) 457nm laser with a power output of 300 mW. Chromomycin A3 fluorescence was collected using a 500nm long pass filter in combination with a 600nm short pass filter. Forward scatter was measured using a (Coherent Sapphire) 488nm laser with a power output of 200mW and this was used as the trigger signal for data collection. Chromosomes were sorted at an event rate of 15000 per second. A 70 micron nozzle tip was used along with a drop drive frequency set to ~96KHz and the sheath pressure was set to 65 PSI. Isolated chromosomes were collected in DNA low-binding tubes containing excess of polyamine buffer.

### Proteomics

FACS sorted mitotic chromosomes were digested with trypsin using an in-Stage Tip digestion protocol^79^ and protein digests were analysed by liquid chromatography-tandem mass spectrometry (LC-MS/MS). Data obtained from triplicate experiments were analysed using the Label-Free Quantification algorithm in MaxQuant^80^ and statistical analysis as well as data visualisation were performed using the Perseus software platform^81^.

### Immunofluorescence

Flow-sorted chromosome (chromosomes 19 and X) were spun onto Poly-L-lysine coated slides (VWR) by cytocentrifugation (Cytospin3, Shandon) at 1,300rpm for 10min, RT. Chromosomes samples were blocked with 6% of normal goat serum for 1h at RT and incubated overnight at 4°C in a humid chamber with the primary antibodies Cenpa (2040S, Cell Signaling), Rad21 (Ab154769, Abcam), Myc (SC40, Santa Cruz), Sox2 (Ab97959, Abcam), Nanog (REC-RCAB0002P-F, 2bScientific), Oct4 (sc-5279, Santa Cruz). Chromosomes where washed (buffer containing 10mM HEPES, 2mM MgCl2, 100mM KCL, and 5mM EGTA) and incubated with appropriate secondary antibodies (anti-mouse-Alexa488 (A11001, Invitrogen), or anti-rabbit-Alexa488 (A11008, Invitrogen), or anti-mouse-A566 (A11031, Invitrogen)) for 1h at RT. Immuno-stained chromosomes were mounted in Vectorshield containing DAPI mounting medium. Super-resolution structured illumination (SIM) microscopy was performed on a Zeiss Elyra microscope using a Plan-Apochromat 63x/1.4 oil objective lens.405nm and 488 nm laser used for fluorescent excitation, and fluorescent emission collected using a bandpass 420-480nm+bandpass, 495-550nm+longpass 650nm filter. Wide-field epi-fluorescence microscopy was performed on an Olympus IX70 inverted microscope using a UPlanApo 100x/1.35 Oil Objective lens.

### TEV Protease cleavage

Total chromosomes (10^7^) or chromosomes 19 (2.10^5^) were incubated with or without the AcTEV protease (Invitrogen) (10 units), 4h at RT with gentle rotation, according to manufacturer’s instruction. Samples were then collected for western blot and prepared for optical imaging and Cryo-Electron Tomography (Cryo-ET).

### Cryo-Electron Tomography

Samples for cryo-ET were prepared by mixing 10μL of flow sorted Chromosomes 19 with 1μL protein-A conjugated to 10nm colloidal gold (CMC Utrecht). 2.5μL of the mixture was pipetted onto freshly glow-discharged Quantifoil Cu/Rh R3.5/1 200 mesh grids and plunge frozen into liquid ethane after removal of excess liquid using a Vitrobot Mark IV (FEI). Frozen grids were transferred to and stored in liquid nitrogen until imaging. Tilt series data was collected on a FEI Titan Krios operating at 300 keV, equipped with a Quantum energy filter and a K2 direct electron detector (Gatan) operating in counting mode, using SerialEM software^82^. Tilt series were collected in two directions starting from 0°, at an unbinned calibrated pixel size of 8.4 Å between ±60° with a 1° increment at 9μm underfocus. A combined dose of 100 e^−^/Å^2^ was applied over the entire series. Tilt series data were aligned and visualised using IMOD^83^.

## Acknowledgements

We would like to thank R. Henderson for for experimental suggestions, C. Tyler-Smith for enabling chromosome sorting, T. Adejumo, F. Pereira, P. Chana for expertise and providing reagents, A. Lisini for reading the manuscript and advice. This work was funded by core support from the medical research council UK to the London Institute of Medical Sciences.

## Author contributions

D.D. and A.G.F. conceived and designed the study. D.D. performed most of the experiments, designed the figures and contributed to writing the manuscript. B.P., H.K., N.V., A.D., A.C.K., A.F., C.W., K.B., and G.Y. conducted experiments and performed analysis. T.A.M.B., A.K.T. performed EM imaging. J.L., J.E., J.G., and B.L.N. provided scientific advice and support. M.M contributed to study design and writing manuscript. A.G.F. conceived and designed the study, wrote the manuscript and supervised the experiments.

