## Supplemental figures and methods for "Systematic identification of factors bound to isolated metaphase ESC chromosomes reveals a role for chromatin repressors in compaction"

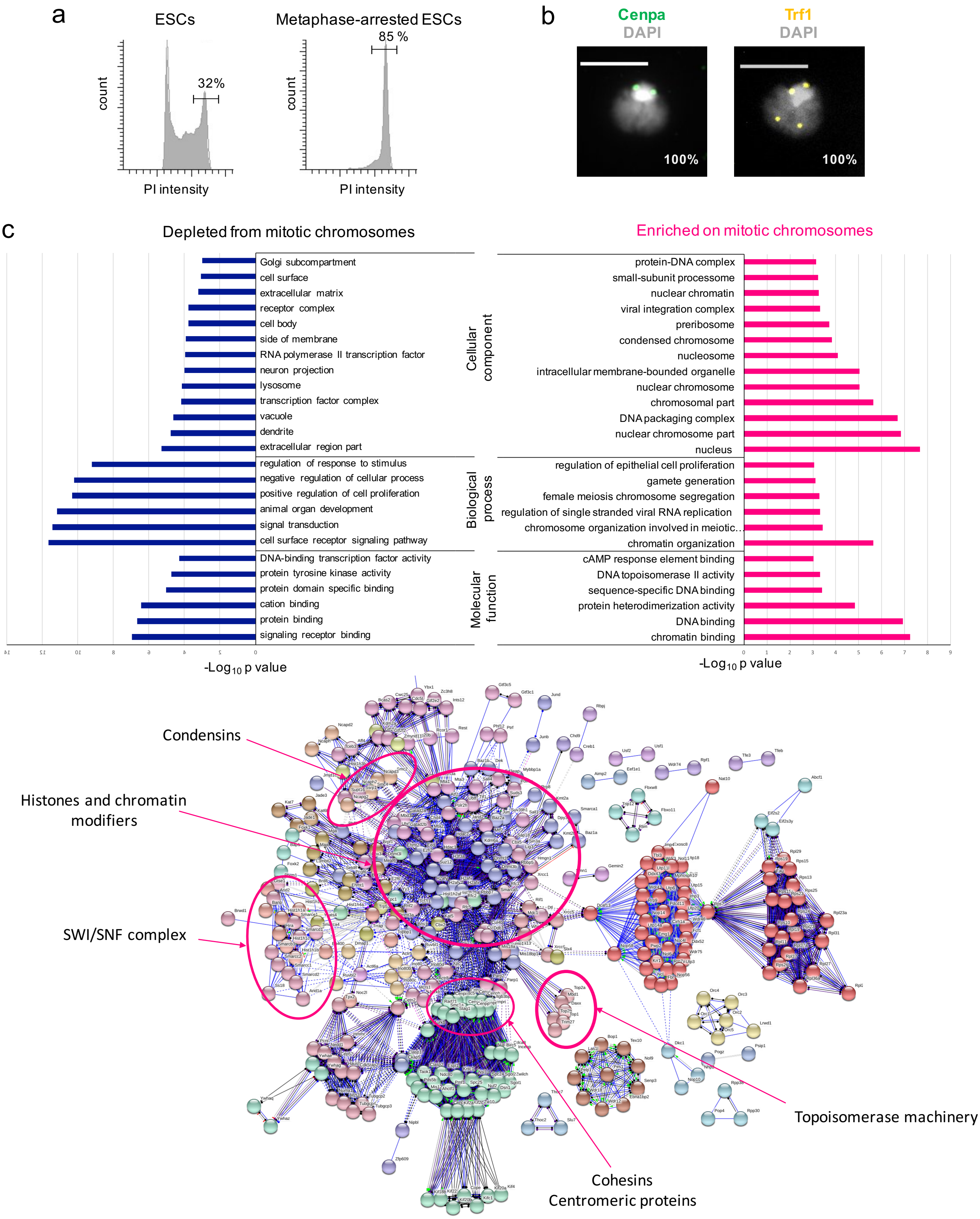

a
ESCs
count
32%
PI intensity
Metaphase-arrested ESCs
85 %
count
PI intensity
b
Cenpa
DAPI
100%
Trf1
DAPI
100%
c
Depleted from mitotic chromosomes
Golgi subcompartment
cell surface
extracellular matrix
receptor complex
cell body
side of membrane
RNA polymerase II transcription factor
neuron projection
lysosome
transcription factor complex
vacuole
dendrite
extracellular region part
regulation of response to stimulus
negative regulation of cellular process
positive regulation of cell proliferation
animal organ development
signal transduction
cell surface receptor signaling pathway
DNA-binding transcription factor activity
protein tyrosine kinase activity
protein domain specific binding
cation binding
protein binding
signaling receptor binding
-Log10 p value
Enriched on mitotic chromosomes
Cellular component
protein-DNA complex
small-subunit processome
nuclear chromatin
viral integration complex
preribosome
condensed chromosome
nucleosome
intracellular membrane-bounded organelle
nuclear chromosome
chromosomal part
DNA packaging complex
nuclear chromosome part
nucleus
Biological process
regulation of epithelial cell proliferation
gamete generation
female meiosis chromosome segregation
regulation of single stranded viral RNA replication
chromosome organization involved in meiotic...
chromatin organization
Molecular function
cAMP response element binding
DNA topoisomerase II activity
sequence-specific DNA binding
protein heterodimerization activity
DNA binding
chromatin binding
0
1
2
3
4
5
6
7
8
9
-Log10 p value
Condensins
Histones and chromatin modifiers
SWI/SNF complex
Topoisomerase machinery
Cohesins
Centromeric proteins

d

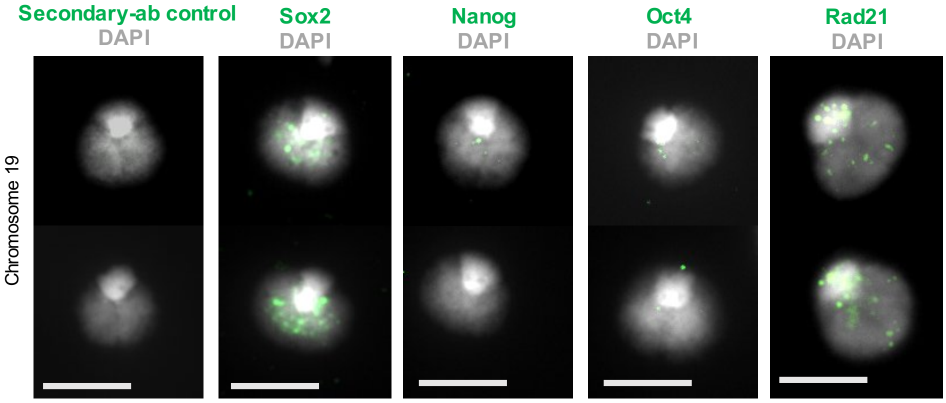


**Supplemental Figure 1**

(a) cell cycle profiles of mouse ESCs determined by staining with propidium iodide (PI intensity), where left panel shows unsynchronised cells and right panel shows samples after 6h treatment with demecolcine. Values indicate percentage cells in G2/M. (b) Representative images of mitotic chromosome 19 isolated from mouse ESCs stained with DAPI (light grey) and labelled to show the distribution of centromeric (CENPA, green) or telomeric (TRF1, yellow) proteins. (c) GO term analysis of proteins identified by LC-MS/MS as significantly enriched (right) or depleted (left) on mitotic ESC chromosomes based on comparisons of sorted chromosomes versus mitotic lysates. Interaction analysis of proteins enriched on mitotic ESC chromosomes is shown as a schematic (lower panel). (d) Representative images of purified ESC mitotic chromosome 19 labelled (green) with antibody to Sox2, Nanog, Oct4 or Rad21 and counterstained with DAPI (light grey). Secondary antibody alone (left) provides a negative control. Scale bars=5 μm.

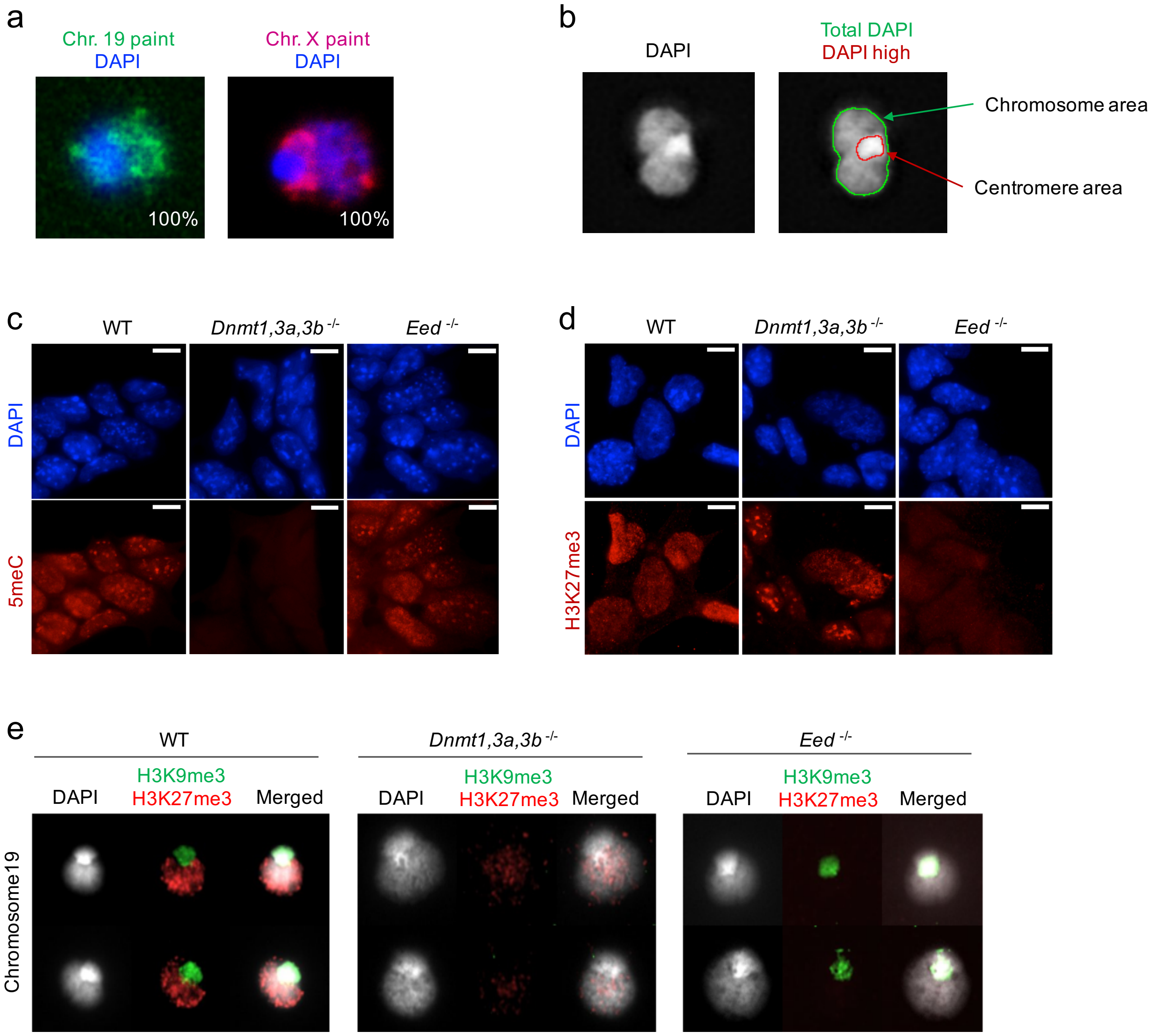


**Supplemental Figure 2**
(a) DNA FISH analysis showing representative images of flow-sorted mitotic chromosomes 19 and X after hybridisation with mouse chromosome 19-specific (green) and X-specific (red) DNA probes, where the percentage values indicate the purity of sorted chromosome samples in each case. (b) Chromosome size measurements were assessed using Fiji/imageJ software to estimate chromosome (total DAPI) and centromere (DAPI high) area, as indicated. (c-d) Relative abundance of 5'methylcytosine (5meC, red) and histone H3K27me3 (red) in asynchronous WT, *Dnmt1,3a,3b*$^{-/-}$ and *Eed*$^{-/-}$ ESCs was assessed by immunofluorescence using DAPI as a counterstain (blue). Scale bars, 8μm. (e) Representative images of immunofluorescence labelling of histone H3K9me3 (green) and H3K27me3 (red) on mouse chromosome 19 from WT, *Dnmt1,3a,3b*$^{-/-}$ and *Eed*$^{-/-}$ ESCs, where DAPI counterstain is shown in light grey.

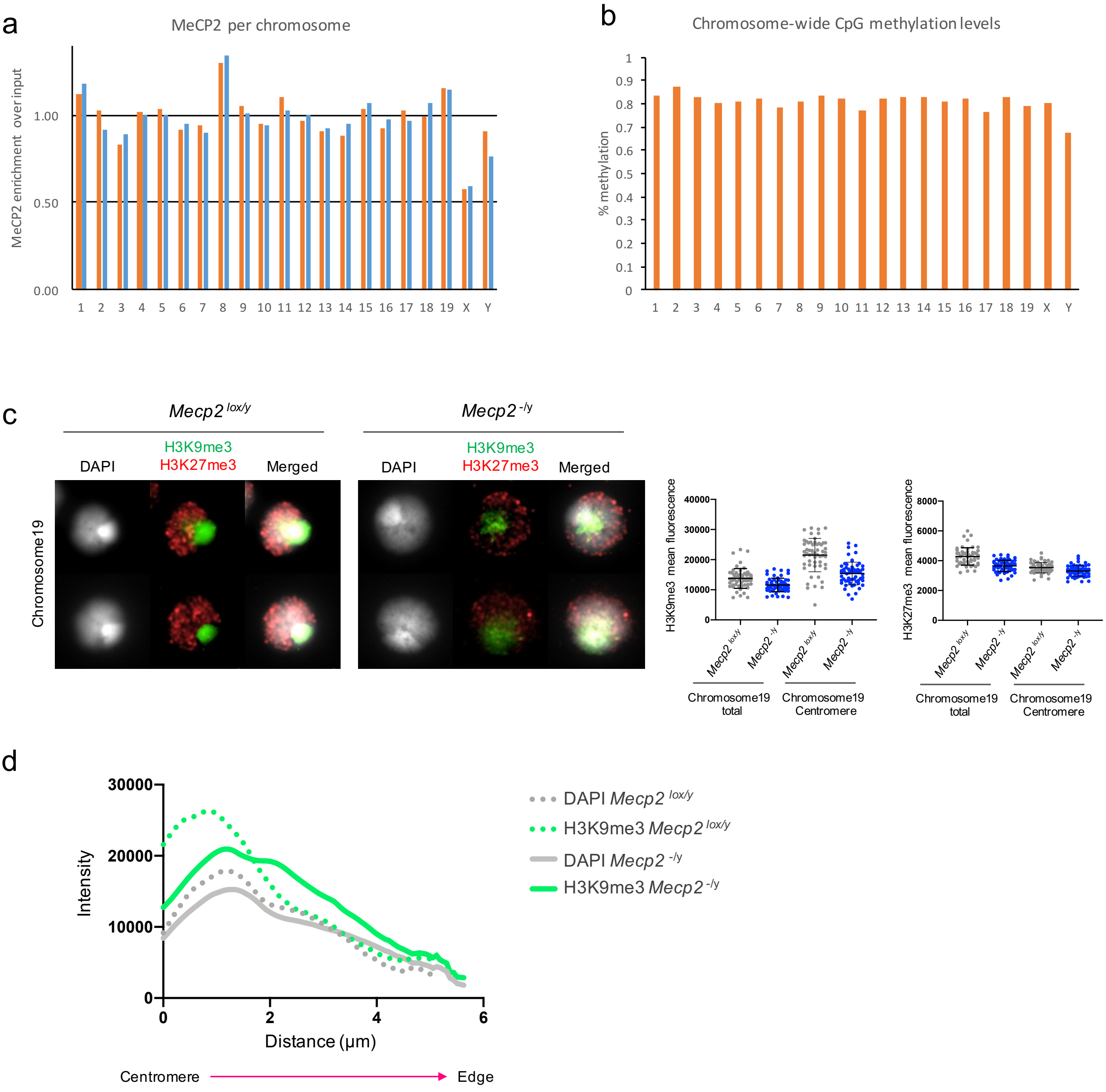


**Supplemental Figure 3**

(a) Biotin-tagged Mecp2 ChIP-seq analysis of male ESCs and post-mitotic neurons, datasets from[1]. Graph shows Mecp2 enrichment (normalized ChIP-seq read count over input) for each chromosomes of ESCs (orange) and mature neuronal cells (blue). (b) Genome-wide methylation analysis of male ESCs, datasets from[2]. Graph show normalized percentage of DNA methylation of each chromosomes. (c) Representative images of immunofluorescence labelling of histone H3K9me3 (green) and H3K27me3 (red) on mouse chromosome 19 from *Mecp2*$^{lox/y}$, *Mecp2*$^{-/y}$ ESCs, where DAPI counterstain is shown in light grey. H3K9me3 (left graph) and H3K27me3 (right graph) mean intensities were measured in both centromere and total chromosome for a minimum of 60 chromosomes for each condition, values indicate mean ± SD. (d) Average distribution plots (minimum of 30 chromosomes) of H3K9me3 and DAPI intensities measured in chromosomes 19 of *Mecp2*$^{lox/y}$ and *Mecp2*$^{-/y}$ ESCs shown in figure 3c.

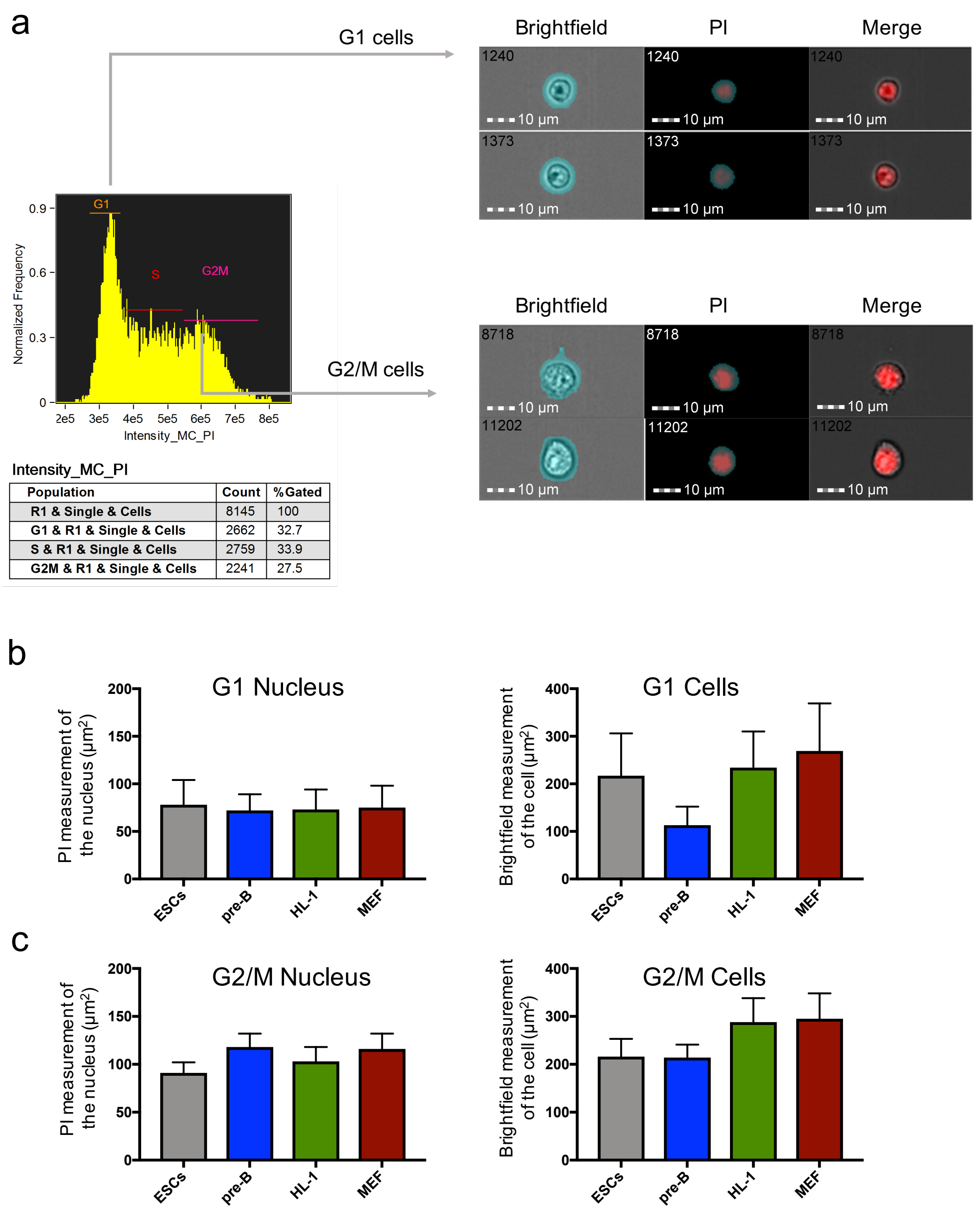


**Supplemental Figure 4**

(a) Nuclear and cellular size measurements of mouse ESCs, pre-B cells, cardiomyocytes (HL-1) and mouse embryo fibroblasts (MEF) were assessed using Amnis image stream (IDEAS software) where cells in G1 or in G2/M were discriminated on the basis of DNA content and PI intensity. Brightfield measurement was used to estimate cell size and refined PI measurements was used to determine nuclear size. (b-c) Histograms summarise the dimensions of nuclei and cells in samples in G1 or in G2/M for each cell type (bars indicate mean+/- SD).

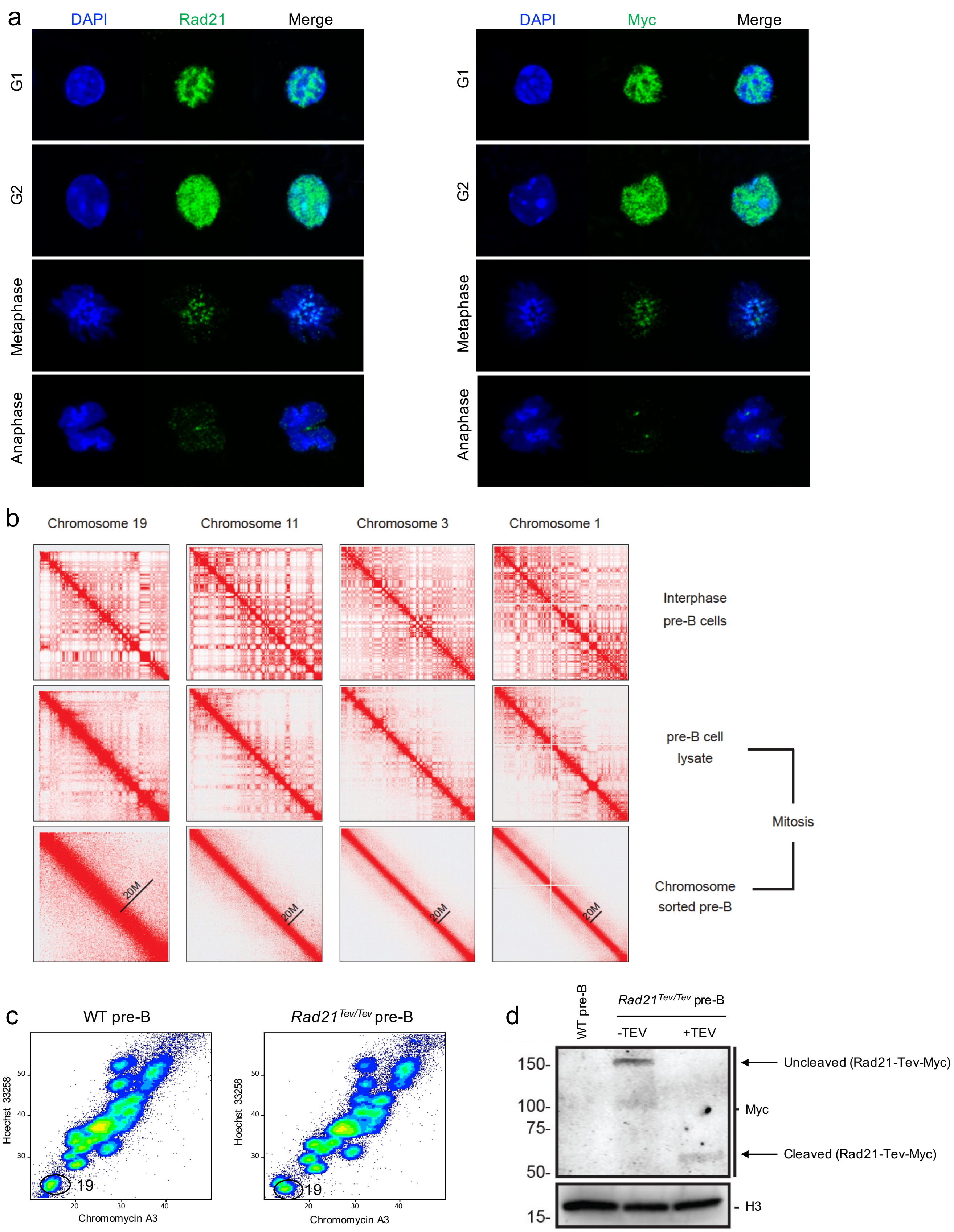


**Supplemental Figure 5**
(a) Immunostaining of Rad21 and Myc in specific cell cycle phases in *Rad21*$^{Tev/Tev}$ pre-B cells, where DAPI stain (blue), Rad21 (green, left panel) and Myc label (green, right panel). (b) Hi-C analysis of flow-sorted chromosomes. Heat-maps displaying contact matrices of chromosome 19, 11, 3, and 1 in interphase pre-B cells (top panel), pre-

B mitotic cell lysates (middle panel) and sorted chromosomes (lower panel), where red colour represents strong interactions. Hi-C data of interphase pre-B cells was downloaded from public repository (GEO accession: GSE82144)[3] (c) Flow karyotype of mitotic chromosomes isolated from WT and *Rad21*$^{Tev/Tev}$ pre-B. Gates used to isolate chromosome 19 are indicated. (d) Immunoblot of Myc-tagged Rad 21 in isolated chromosomes of WT and *Rad21*$^{Tev/Tev}$ pre-B cells after TEV protease cleavage. Arrows indicate the uncleaved Rad21 (top) and the Rad21-cleaved fragment (lower arrow). Histone H3 was used as a loading control for the western blot.

### Supplemental methods

#### Proteomics

Flow-sorted mitotic chromosomes were pelleted by centrifugation (20,000g, 10min, 5°C). Supernatants were removed and the obtained pellet was processed by in-Stage Tip digestion[4] using commercially available iST tips (Preomics, Martinsried, Germany) according to manufacturers recommendations. Briefly, pellets were suspended in lysis buffer and heat denatured, reduced and alkylated on a heated shaking incubator (1,000rpm, 10min, 95°C). DNA was fragmented by sonication on an ultrasonic water bath (10min) and samples were digested with trypsin (500rpm, 1h, 37°C). Sample clean-up and desalting was carried out in the iST device using the recommended wash buffers. Peptides were eluted with elution buffer (2x100ul), concentrated in a centrifugal evaporator and resuspended in LC loading buffer (20ul). LC-MS/MS analysis was performed as follows. Resuspended protein digests were transferred to auto sampler vials for LC-MS analysis. Peptides were separated using an ultimate 3000 RSLC nano liquid chromatography system (Thermo Scientific) coupled to a Q-Exactive HF-X tandem mass spectrometer (Thermo Scientific) via an EASY-Spray source. Sample volumes were loaded onto a trap column (Acclaim PepMap 100C18, 100umx2cm) at 8ul/min in 2% acetonitrile, 0.1% TFA. Peptides were eluted on-line to an analytical column (EASY-Spray PepMap C18, 75um x 75cm). Peptides were separated at 200nl/min using a ramped 120min gradient from 1-42% buffer B (buffer A: 5% DMSO, 0.1% formic acid; buffer B: 75% acetonitrile, 0.1% formic acid, 5% DMSO). Eluted peptides were analysed operating in positive polarity using a data-dependent acquisition mode. Ions for fragmentation were determined from an initial MS1 survey scan at 120,000 resolution (at m/z 200) in the Orbitrap followed by HCD (Higher-energy Collisional Dissociation) of the top 30 most abundant. MS1 and MS2 scan AGC targets set to 3e6 and 5e4 for a maximum injection time of 25ms and 50ms, respectively. A survey scan covering the range of 350 –1750m/z was used, with HCD parameters of isolation width 1.6m/z and a normalised collision energy of 27%. Data obtained from biological triplicate experiments (each loaded in duplicates) were analysed using the Label-Free Quantification algorithm in MaxQuant software platform (v1.6.2.3)[5] with database searches carried out by the in-built Andromeda search engine against the Swissprot Mus musculus database (16,950 entries, v.20180104). A reverse decoy database was created and results displayed at a 1% false-discovery rate (FDR) for peptide spectrum matches and protein identifications. Search parameters included: trypsin, two missed cleavages, fixed modification of cysteine carbamidomethylation and variable modifications of methionine oxidation, asparagine deamidation and protein N-terminal acetylation. Label-free quantification was enabled with an LFQ minimum ratio count of 2. 'Match between runs' function was used with match and alignment time limits of 0.7 and 20 min, respectively. Statistical analysis as well as data visualisation were performed using the Perseus software platform[6].

**Nuclear size measurements**

Cells were labelled with Propidium iodide as describes. Data acquisition was performed on Amnis image stream flow cytometer and analyzed using Amnis IDEAS software.

**Telomeres labeling**

ESCs were transfected with 12ug of TRF1-YFP plasmid[7]. TRF1-YFP positive cells were sorted using BD AriaIII flow sorter and grown in normal ESCs medium. Metaphase chromosomes 19 and X from TRF1-YFP positive cells were sorted and analyzed by optical imaging.

**Chromosome painting**

Flow-sorted chromosomes 19 and X ($10^5$) were spun onto Poly-L-lysine coated slides by cytocentrifugation (Cytospin3, Shandon) at 1,300rpm for 10min, RT. Fluorescent *in situ* hybridization was performed as previously described (Geigl et al 2006). Samples were hybridized with mouse chromosomes 19 (D-1419-050-FI) ox X paints (D-1420-050-OR) according to manufacturer's instruction.

**Hi-C on isolated chromosomes**

Hi-C was performed as previously described[8]. Briefly, sorted chromosomes were cross-linked in 1% formaldehyde for 10min at RT. Chromatin was digested with 600 units of HindIII overnight at 37°C with rotation. Digested sticky ends were filled in by DNA polymerase I, large Klenow fragment in the presence of 50nM biotin-14-dATP, 50nM dTTP, 50nM dGTP, and 50nM dCTP for 90min at 37°C with rotation. Chromatin fragment ends were ligated by 4000 units of T4 DNA ligase by incubating at RT for 6h with slow rotation. Proteins were digested by proteinase K and then cross-linked chromatin was reversed by incubating overnight. RNase A was used to remove RNA after decross-linking. After isolation with phenol / chloroform / isoamyl alcohol (25:24:1 mixture), DNA was precipitated by sodium acetate / ethanol precipitation. DNA was sheared for 9min by Bioruptor sonicator (30sec on and 30sec off per minute using high power setting). DNA fragments in the range of 300-500bp were selected with AMPure XP beads. After biotinylated DNA was captured on Dynabeads MyOne Streptavidin T1, DNA ends were repaired in a mixture of T4 polynucleotide kinase, T4 DNA polymerase I and DNA polymerase I, large Klenow fragment. dATP was then added to the repaired ends using Klenow Fragment (3'→5' exo-). NEBNext adaptors for Illumina were ligated to the dA-tailed ends. After USER enzyme digestion, NEBNext oligos for Illumina were used for library preparation. A PCR titration was performed to determine the minimal number of PCR cycles (8 cycles in this work). Hi-C libraries were sequenced on an Illumina HiSeq 2500 sequencer. 2×100 bp paired-end reads were generated for downstream analysis. Hi-C data was mapped and processed using bowtie 2 (v2.3.2) and HiC-Pro (v2.7.8) by default setting[9, 10]. Raw sequencing data was mapped to Mus musculus genome (UCSC assembly mm9, NCBI build 37). PCR duplicates and read pairs that aligned on the same restriction fragment were removed. After converting using the HiC-Pro hicpro2juicebox.sh, valid chromatin contacts were normalized by the KR matrix balancing algorithm and hic files were created using the Juicer Pre[11]. Heat-maps of chromatin interaction matrix was visualized and generated using the Juicebox[12]. To plot relative frequency of intrachromosomal interactions, genomic distances from 50 kb to 100 Mb were divided into logarithmically-increasing bins in size. The interaction separations measured were started from 50 kb and then increased with a factor 1.12. Average values of chromatin interactions were calculated for each separation. Contact frequency of these genomic distances was normalized to the highest point and then plotted with a log-log transformation after smoothing using the ggplot R package[13].
